# Co-Expressed MicroRNAs Identify Potential Mechanisms Underlying Risk for Multimorbid Depression and Type 2 Diabetes in Midlife Women

**DOI:** 10.64898/2026.06.10.731370

**Authors:** Kayla D. Longoria, Benjamin M. Stroebel, Meghana Gadgil, Sandra Weiss, Kimberly A. Lewis, Nicole Perez, Elena Flowers

## Abstract

**Background:** Women are disproportionately affected by multimorbid depression and type 2 diabetes (T2D), with prevalence peaking during midlife (40-64 years), a biologically dynamic timeframe due to changes associated with reproductive aging. Yet, phenotypic and mechanistic factors contributing to midlife women’s disproportionate risk for co-occurrence remain poorly defined. We previously identified co-expressed microRNAs (miRs) in midlife women with prediabetes that increased odds of assignment to a high psychometabolic risk phenotype. Here, we extend these findings by characterizing putative mRNA targets of these co-expressed miRs and pathways overrepresented among mRNAs, providing insights into potential mechanisms underlying psychometabolic risk in midlife women.

**Methods:** This study included baseline data from midlife women (ages 40-64 years) with prediabetes who participated in the Diabetes Prevention Program (DPP) (n = 603). *In silico* analyses were performed using miRTarBase to identify mRNAs regulated by 3 or more of the miRs that most prominently loaded a principal component previously identified to increase odds of assignment to a high psychometabolic risk phenotype defined in this sample. Pathway enrichment analysis was conducted to assess for overrepresentation of KEGG pathways among predicted mRNA targets. To enhance interpretability, pathways were thematically clustered based on their evidenced role in human physiology.

**Results:** We identified a total of 13 mRNAs targeted by co-expressed miRs associated with increased odds of assignment to a high psychometabolic risk phenotype in midlife women with prediabetes. Pathway enrichment analysis revealed a total of 71 KEGG pathways with overrepresentation of identified mRNA targets. Four overarching biological themes emerged, reflecting involvement of metabolic, inflammatory, endocrine, and stress/biological weathering-related processes.

**Conclusions:** Experimentally validated mRNA targets related biological pathways were identified, providing multisystem insights into potential mechanisms underlying risk for multimorbid depression and T2D in midlife women. Findings offer mechanistic targets for experimental validation and future precision health research focused on this high-risk population. Overall, this work positions the utility of miRs as context-sensitive biomarkers in the characterization of risk for complex, multimorbid conditions in women during biologically dynamic timeframes.

## BACKGROUND

Depression and Type 2 Diabetes (T2D) are leading contributors of global disability with rising incidence rates and earlier age of onset [1]. While each condition is independently associated with poor clinical outcomes, multimorbid depression and T2D compound risk for accelerated disease progression and greater morbidity [2–4]. Women are disproportionately affected by this multimorbidity dyad, and prevalence peaks during midlife (ages 40-64 years) [5–7]. For women (female sex) with intact ovarian function, midlife represents a biologically distinct timeframe, as physiological changes associated with reproductive aging (pre-, peri-, post-menopause) gradually occur over the course of multiple years [8]. The Study of Women’s Health Across the Nation (SWAN), one of the most comprehensive longitudinal studies on women’s health during midlife in the U.S., indicates midlife is a high-risk period for mood disturbances (e.g., depression symptoms) and metabolic shifts known to increase risk for T2D (e.g., visceral adiposity, dyslipidemia, insulin resistance) [8–11]. Despite this, few sex- and age-specific investigations have examined phenotypic and mechanistic drivers underlying women’s risk for multimorbid depression and T2D during midlife, limiting advances in effective strategies for earlier detection and intervention.

Depression and T2D have a bidirectional relationship in which one condition increases risk for the other [4,5,12]. Few studies, however, have integratively examined depression symptoms and metabolic risk factors to comprehensively characterize psychometabolic risk phenotypes, with those focused on midlife women being particularly rare. Consequently, mechanisms contributing to risk for co-occurrence remain incompletely understood, with current knowledge largely limited to evidence derived from independent investigations focused on single systems defined *a priori*, and, when examined concurrently, is often studied within the context of a diagnosis or comorbidity [13,14]. Because women are disproportionately affected by this multimorbidity dyad during a timeframe that overlaps with physiological changes associated with reproductive aging [8], a systems-level approach is needed to delineate biological processes contributing to risk for co-occurrence within the unique context of this biological milieu. Although representation of female sex in biomedical research is improving, critical knowledge gaps on mechanisms contributing to risk for complex, multimorbid conditions in women during key reproductive life stages remain under-studied [15].

MicroRNAs (miRs) are post-transcriptional regulators that bind to messenger RNAs (mRNAs) to modulate gene expression, with individual miRs often co-regulating mRNA targets to coordinate physiological processes across multiple biological systems. Circulating miRs measured in blood exhibit dynamic changes in response to external exposures; thus, can provide insights into adaptive and maladaptive responses in the context of human health and disease [16,17]. Because miRs have the potential to integrate signals across domains (e.g., biological, environmental) and reflect temporal changes, they are emerging as promising biomarkers for personalized screening and intervention of complex conditions, including T2D and depression [18–21]. However, limited research has leveraged co-expressed miRs anchored to a multimorbid risk phenotype to advance a more comprehensive understanding of mechanisms underlying risk for depression and T2D co-occurrence, especially in midlife women.

Our prior study in a sample of midlife women with prediabetes showed that elevated risk for both depression and T2D clustered together in what was defined as a high psychometabolic risk phenotype [22]. We also identified a cluster of co-expressed miRs that increased odds of assignment to this high-risk phenotype. To bridge critical gaps in understanding of mechanisms underlying risk for depression and T2D co-occurrence in midlife women, this paper will characterize mRNA targets and related biological pathways that are regulated by the co-expressed miRs that are associated with the high-risk phenotype. This study will provide multisystem insights into potential mechanisms underlying risk for multimorbid depression and T2D in midlife women, offering mechanistic targets for further investigation in this high-risk population.

## METHODS

This secondary analysis leveraged baseline data from midlife (ages 40-64 years) women with prediabetes (n = 603) who participated in the Diabetes Prevention Program (DPP) trial. Details of the parent DPP trial have been described in detail previously [23–25].

Previously, to identify how depression symptoms and metabolic risk factors phenotypically cluster in midlife women, our prior secondary analysis in this sample performed unsupervised k-means clustering integrating baseline clinical characteristics commonly evaluated independently when screening for T2D and depression in primary care settings [22]. Variables included: age, glycemic biomarkers (i.e., fasting blood glucose (FBG), hemoglobin A1c (HbA1c)), anthropometric measures of adiposity (i.e., body mass index (BMI), waist circumference), lipid measures (i.e., high-density lipoprotein (HDL), triglycerides), and depression symptom scores measured by the Beck Depression Inventory (BDI-I) that was administered in the parent DPP trial [24,26]. Two psychometabolic risk phenotypes emerged (i.e., high vs. low) and in this sample of women with prediabetes, both phenotypes demonstrated elevated risk for T2D. However, the high-risk phenotype exhibited significantly greater risk for T2D across all metabolic features except triglycerides and significantly higher depressive symptom scores, reflecting elevated risk for multimorbid depression and T2D compared to the low-risk phenotype characterized by T2D risk among two metabolic domains (i.e., glycemic biomarkers, measures of adiposity).

Principal component analysis (PCA) was performed to reduce dimensionality of miR measures and derive uncorrelated PCs representing shared variation in miR expression. Logistic regression models including all four miR PCs were applied to determine PCs that were significantly associated with the high-risk phenotype, adjusting for race and ethnicity. One PC consistently and significantly showed an increased association with the high-risk phenotype, with the 10 miRs with the highest loadings onto this PC forming the basis for the *in silico* analyses described in this manuscript.

### Molecular Data Collection

The Fireplex Multiplex Circulating MicroRNA Assay (Abcam, MA) was used for direct quantification of 58 miRs from plasma collected at the baseline visit of the DPP trial [19]. MiRs were hybridized to complementary oligonucleotides covalently attached to encoded hydrogel microparticles. The bound target was ligated to oligonucleotide adapter sequences that serve as universal PCR priming sites. The miR-adapter hybrid models were then denatured from the particles and reverse transcription polymerase chain reaction (RT-PCR) was performed using a fluorescent forward primer. Once amplified, the fluorescent target was rehybridized to the original capture particles and scanned on an EMD Millipore Guava 6HT flow cytometer (Merck KGaADarmstadt, Germany).

### Statistical Analysis

*In silico* analyses were performed to determine predicted mRNAs and biological pathways regulated by the 10 miRs that most prominently loaded (loading coefficient >0.2) onto the PC that was significantly associated with the high-risk phenotype. These included miR-320a, miR-320c, miR-126, miR-486, miR-93, miR-17, let-7c, let-7f, miR-30a, and miR-122 [22]. based on our prior review of tools for miR target prediction [19,28], we used MiRTarBase (version 10.0) [27] to identify predicted mRNA targets of these 10 miRs. The *Homo sapiens* microRNA-target interaction (MTI) database was downloaded from miRTarBase, and R [29] was used to filter mRNA targets of the identified miRs to include only those with strong, experimentally validated evidence (i.e. Western blot, qRT-PCR, and reporter). Within OrganismDbi v1.36.0, the Homo.sapiens package (v1.3.1) was used to determine the entrez IDs for target mRNAs, which were added to the miRTarBase results. Pathway enrichment analysis was conducted to identify Kyoto Encyclopedia of Genes and Genomes (KEGG) [30] pathways with overrepresentation of mRNA that are regulated by the identified miRs using the clusterprofiler package v4.2.2. Significance was determined by a false-discovery rate adjusted p-value <0.05 [31]. To determine broad biological domains involved in risk for multimorbid depression and T2D in midlife women, pathways not largely limited to cancer or infectious disease were organized into biological themes based on evidence linking dysregulation to disease pathology (i.e., depression, T2D) in human populations.

## RESULTS

The subset of midlife women with prediabetes from the DPP trial who were included in this study had a mean age of 50 ± 9 years, with 59% categorized as White race and 15% Hispanic ethnicity. Overall, anthropometric and clinical measures indicated this sample was obese (34.9 ± 7.3kg/m^2^) and had elevated waist circumference (103 ± 7.3cm), FBG (106 ± 7mg/dL), HbA1c (5.9 ± 0.5%), and triglycerides (161 ± 90mg/dL), and low HDL (49 ± 13mg/dL). The mean depressive symptom score was 4.9 ± 4.4, falling within the category of minimal depressive symptoms [26]. The high-risk phenotype, which was younger in chronological age (48 ± 9 years) and had a higher proportion of midlife women racially categorized as Black (36%), exhibited significantly higher risk for T2D across all metabolic risk factors except triglycerides, as well as significantly higher depressive symptom scores (6.6 ± 5.2) (**Table 1**).

**Table 1.** Demographic and Clinical Characteristics by Cluster.

| **Characteristic**<br>**mean ± SD or n (%)** | **Clusters** |  |  | **p-value** |
| --- | --- | --- | --- | --- |
|  | **Overall**<br>**(n=603)** | **Low Risk**<br>**(n=361)** | **High Risk**<br>**(n=242)** |  |
| **Age (years)** | 50 ± 9 | 51 ± 10 | 48 ± 9 | <0.0001 |
| **BMI kg/m<sup>2</sup>)** | 34.9 ± 7.3 | 30.6 ± 3.9 | 41.3 ± 6.4 | <0.0001 |
| **Waist (cm)** | 103 ± 15 | 94 ± 9 | 117 ± 12 | <0.0001 |
| **FBG (mg/dL)** | 106 ± 7 | 104 ± 6 | 110 ± 8 | <0.0001 |
| **HbA1c (%)** | 5.9 ± 0.5 | 5.8 ± 0.4 | 6.1 ± 0.5 | <0.0001 |
| **Triglycerides (mg/dL)** | 161 ± 90 | 159 ± 92 | 163 ± 86 | 0.5272 |
| **HDL (md/dL)** | 49 ± 13 | 52 ± 13 | 44 ± 10 | <0.0001 |
| **BDI depression score** | 4.9 ± 4.4 | 3.9 ± 3.4 | 6.6 ± 5.2 | <0.0001 |
| **Race and Ethnicity** |  |  |  | 0.0004 |
| **White** | 357 (59) | 234 (65) | 123 (51) |  |
| **Black** | 138 (23) | 52 (14) | 86 (36) |  |
| **Hispanic** | 88 (15) | 58 (16) | 30 (12) |  |
| **Other** | 20 (3) | 17 (5) | 3 (1) |  |

SD – standard deviation; BMI – body mass index; FBG – fasting blood glucose; HbA1c – hemoglobin A1c; HDL – high density lipoprotein cholesterol; BDI – Beck Depression Inventory;

A total of 13 predicted mRNAs were identified to be targeted by three or more of the miRs with the highest loading values on the PC that was significantly associated with the high psychometabolic risk phenotype (**Table 2**). Pathway enrichment analysis revealed a total of 71 KEGG pathways with statistically significant overrepresentation of the 13 identified mRNAs, with 31 that are not primarily associated with cancer or infectious disease (**Supplementary Table 1, Supplementary Table 2**). From these pathways, four overarching biological themes emerged: metabolic, inflammatory, endocrine, and stress/biological weathering (**Table 2**). The mRNAs differed in their pathway distribution, with some predominantly represented within specific themes whereas others are located in pathways more evenly distributed across all four themes (**Figure 1**).

**Table 2.** Genes and Associated KEGG Pathways Regulated by Three or More MiRs Associated with Elevated Psychometabolic Risk.

| miRs | GeneID | KEGG Pathway Themes |  |  |  |  |
| --- | --- | --- | --- | --- | --- | --- |
|  |  | Total Pathways<br>(n) | Metabolic<br>(n) | Inflammatory<br>(n) | Endocrine<br>(n) | Stress/Biol<br>Weather<br>(n) |
| 320a, 93, 17, 7c, 7f | *MYC* | 11 | 1 | 3 | 2 | 5 |
| 486, 17, 7c, 7f | *CCND1* | 15 | 5 | 1 | 3 | 6 |
| 486, 93, 17, 122 | *TGFB1* | 12 | 3 | 5 | 1 | 3 |
| 93, 17, 30a | *CDH1* | 4 | 1 | 0 | 0 | 3 |
| 93, 17, 7f | *CDKN1A* | 10 | 2 | 1 | 3 | 4 |
| 126, 93, 17 | *E2F1* | 4 | 0 | 0 | 2 | 2 |
| 93, 17, 30a | *HIF1A* | 5 | 0 | 2 | 1 | 2 |
| 320a, 7c, 122 | *IGF1R* | 12 | 5 | 1 | 1 | 5 |
| 126, 17, 30a | *JAK1* | 5 | 1 | 3 | 0 | 1 |
| 320a, 93, 17 | *PTEN* | 7 | 3 | 0 | 0 | 4 |
| 486, 17, 122 | *SMAD3* | 13 | 5 | 2 | 0 | 6 |
| 320a, 93, 17 | *TGFBR2* | 12 | 4 | 4 | 1 | 3 |
| 126, 93, 17 | *VEGFA* | 8 | 2 | 1 | 1 | 4 |
| **Total pathways** |  | 30 | 8 | 6 | 5 | 11 |

*CCND1* - Cyclin D1; *CDH1* - Cadherin 1; *CDKN1A* - Cyclin Dependent Kinase Inhibitor 1A; *E2F1* - E2F Transcription Factor 1; *HIF1A* - Hypoxia Inducible Factor 1 Subunit Alpha; *IGF1R* - Insulin Like Growth Factor 1 Receptor; *JAK1* - Janus Kinase 1; KEGG – Kyoto Encyclopedia of Genes and Genomes; *MYC* - MYC Proto-Oncogene, BHLH Transcription Factor; *PTEN* - Phosphatase and Tensin Homolog; *SMAD3* - SMAD Family Member 3; *TGFB1* - Transforming Growth Factor Beta 1; *TGFBR2* - Transforming Growth Factor Beta Receptor 2; *VEGFA* - Vascular Endothelial Growth Factor A

**Figure 1.**
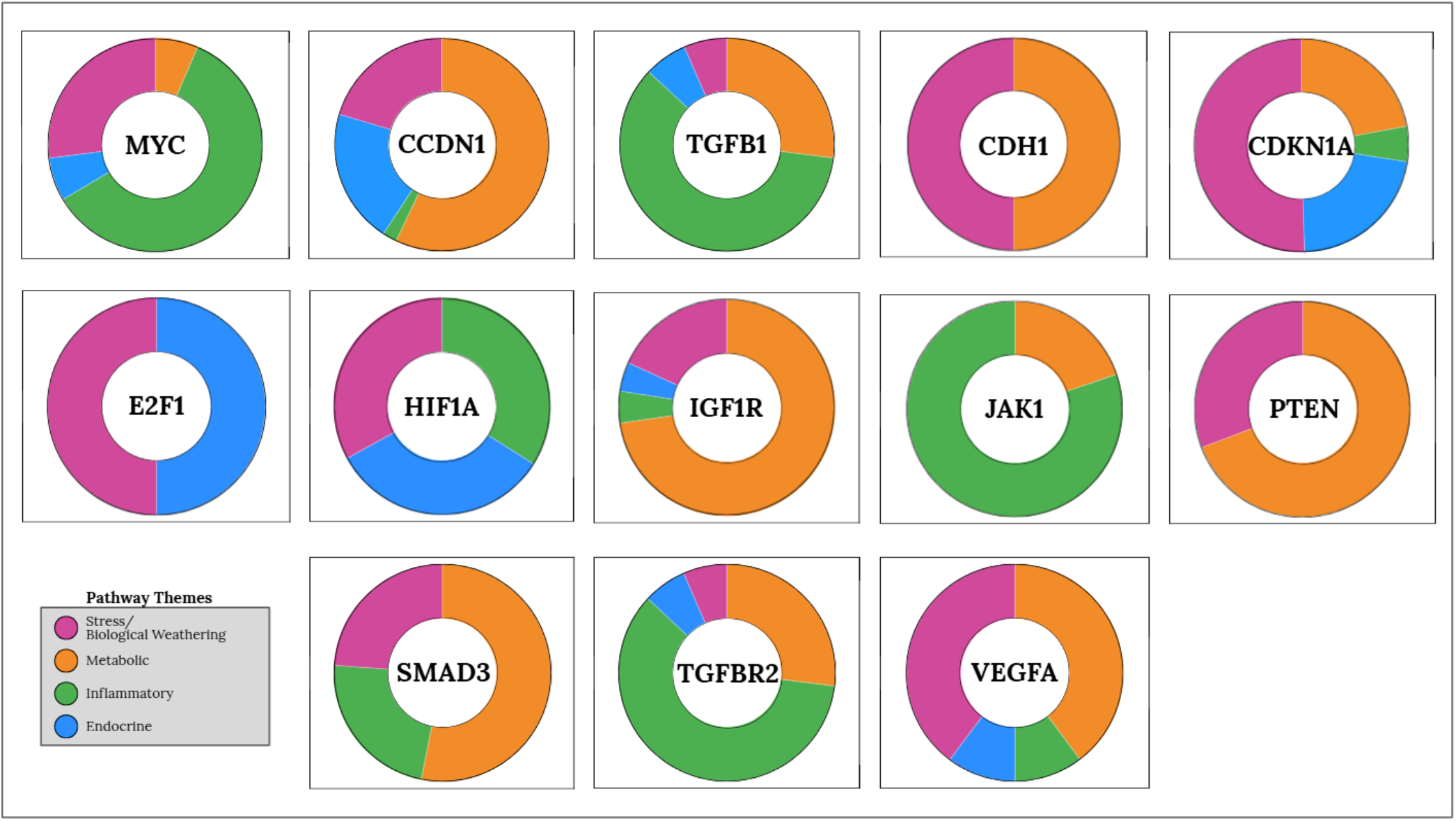
mRNAs Targeted by MicroRNAs Associated with Elevated Psychometabolic Risk and Their Related Biological Pathways that are Related to Depression and/or Type 2 Diabetes **Legend**. White circles show mRNA name. Outer rings shows the mechanistic theme. *CCND1* - Cyclin D1; *CDH1* - Cadherin 1; *CDKN1A* - Cyclin Dependent Kinase Inhibitor 1A; *E2F1* - E2F Transcription Factor 1; *HIF1A* - Hypoxia Inducible Factor 1 Subunit Alpha; *IGF1R* - Insulin Like Growth Factor 1 Receptor; *JAK1* - Janus Kinase 1; KEGG – Kyoto Encyclopedia

Of the 31 pathways related to depression or T2D pathology, a subset of these pathways are potentially related to both conditions independently. In the theme of metabolic activity, pathways included P13K-AKT signaling, AGE-RAGE signaling pathway in diabetic complications, AMP-activated protein kinase (AMPK), Forkhead box O (FoxO), Apelin signaling, and mechanistic Target of Rapamycin (mTOR) signaling. Pathways related to inflammatory processes that are potentially related to both T2D and depression included Janus kinase/signal transducers and activators of transcription (JAK-STAT) signaling, transforming growth factor-beta (TGF-*β*), Mitogen-activated protein kinase (MAPK) signaling, Efferocytosis, and Th17 cell differentiation. Within the theme of endocrine activity, pathways included Oxytocin signaling, Thyroid hormone signaling, Endocrine resistance, and Relaxin signaling. Pathways within the theme of stress/biological weathering processes included Cellular senescence, Wnt signaling, Hypoxia-inducible factor 1 (HIF), Rap1 signaling, Hippo signaling, and p53 signaling (**Table 2**). **Figure 2** illustrates relationships between miRs and their mRNA targets (Panel A), mRNAs and their enriched pathways (Panel B) and how these pathways flow into respective biological themes (Panel C).

**Figure 2.**
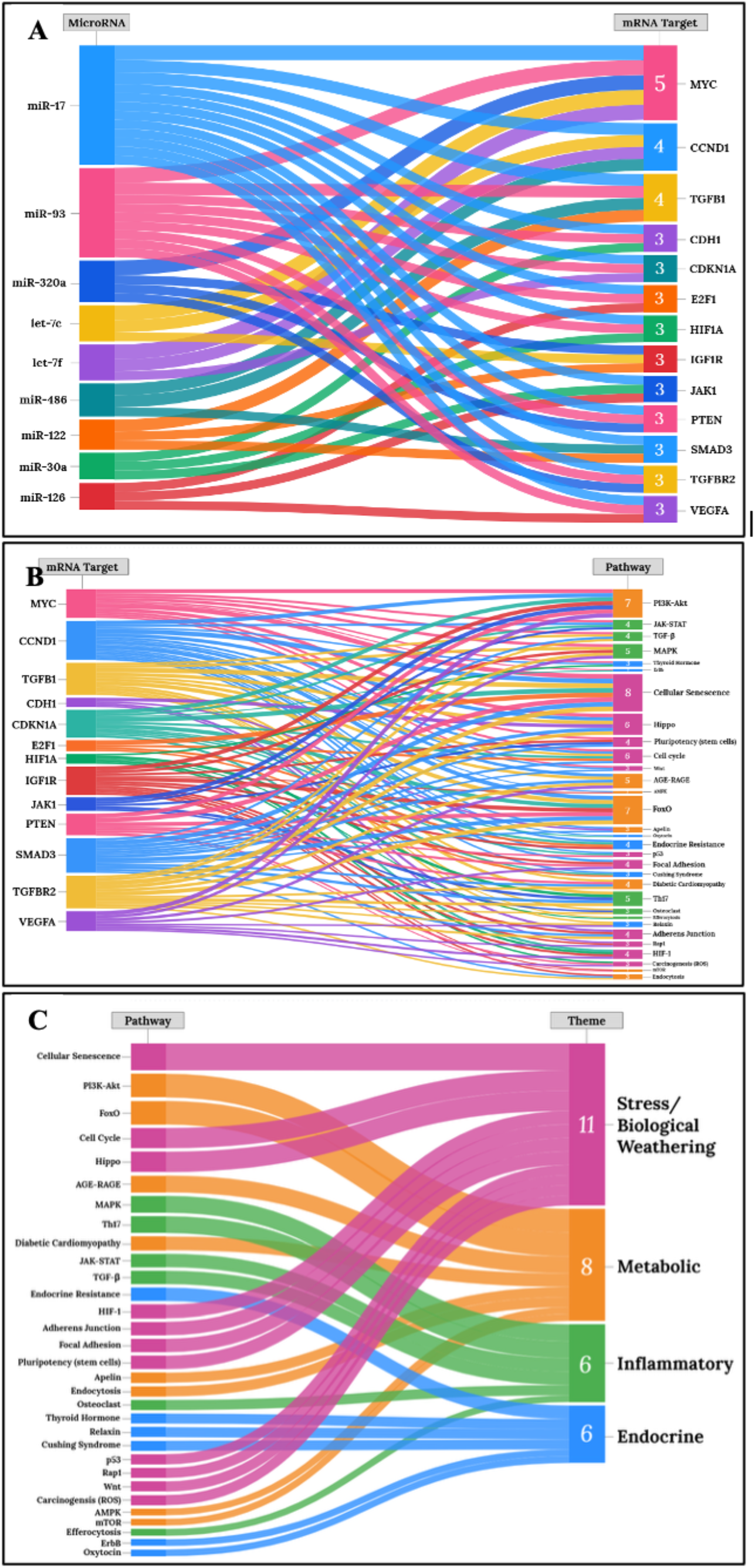
Sankey Plots Showing Relationships Between miRs Associated with Elevated Psychometabolic Risk and their mRNA Targets and Associated KEGG Pathways **Legend**. Panel A shows mRNAs that are targeted by the set of miRs associated with elevated psychometabolic risk. Panel B shows the KEGG pathways that contain overrepresentation of the mRNAs from Panel A. Panel C shows these KEGG pathways organized by mechanistic them. KEGG – Kyoto Encyclopedia of Genes and Genomes

## DISCUSSION

This study extends our prior study that showed co-expressed miRs are associated with increased psychometabolic risk in midlife women by identifying putative mRNA targets of these miRs and their related pathways. These findings broadly support observations that depression and T2D commonly co-occur due to an overlap in behavioral and biological factors and provide systems-level insights into candidate mechanisms that may signal risk for co-occurrence in midlife women. Overall, these findings validate multisystem involvement in women’s risk for depression-T2D co-occurrence and support the utility of miRs as comprehensive context-sensitive biomarkers for women’s risk for complex, multimorbid conditions during biologically dynamic timeframes (e.g., pre-, peri-, post-menopause associated changes [midlife]).

Cellular senescence, a hallmark of biological aging [32], emerged as the most consistently enriched pathway across mRNA targets. Senescence is a homeostatic response to stress (e.g., telomere shortening, environmental exposures) in which damaged or overly stressed cells experience irreversible cell cycle arrest to prevent proliferation but remain biologically active, secreting a complex array of molecules (e.g., interleukin-8 (IL-8), transforming growth factor-beta (TGF-β)) known as the senescence-associated secretory phenotype (SASP) [32–34]. Although initially protective, chronic senescence activation due to prolonged or repeated stress exposures (e.g., environmental, psychosocial, behavioral) can overwhelm immune-mediated clearance, leading to senescent cell accumulation beyond what is expected for chronological age, reflecting biological age acceleration [32]. This pattern aligns with allostatic load and biological weathering theories, which posit that prolonged exposures to stress become biologically embedded through multisystem dysregulation, contributing to disparities in chronic disease outcomes [35–38]. The prominent enrichment of cellular senescence across mRNA targets, coupled with significant phenotypic differences observed in the high-risk group (i.e., younger chronological age, higher proportion of Black race, elevated T2D risk and depressive symptoms), aligns with evidence that broadly supports these theories [35,37,39–41]. Given the well-documented relationship between chronic stress and clinical and subclinical depressive symptoms [42], which has impacts for health behaviors including those related to T2D prevention and management [43], these findings suggest that chronic stress may be a mechanism for psychometabolic multimorbidity [44]. The DPP trial did not directly measure chronic stress exposure and future longitudinal studies incorporating stress measures, including environmental and psychosocial, can clarify how biopsychosocial and behavioral interactions shape women’s risk for multimorbid depression and T2D in midlife.

While senescence is considered a multisystem process involving interconnected pathways that span multiple biological systems, SASP factors (e.g., pro-inflammatory cytokines, chemokines, growth factors, proteases) are among the most widely recognized features of senescence biology [33]. In addition to enrichment of cellular senescence, multiple pathways known to modulate SASP induction or maintenance were also prominently represented in the present study (i.e., Mitogen-activated protein kinase (MAPK), Janus kinase-signal transducer and activator of transcription (JAK-STAT), TGF-β; Mechanistic target of rapamycin (mTOR), and Tumor protein 53 (p53)). Additionally, the Transforming Growth Factor Beta 1 (*TGFB1*) mRNA, a well-established component of SASP [45], was targeted by four of the co-expressed miRs that are associated with the high psychometabolic risk phenotype. *TGFB1* is considered a pleiotropic cytokine, meaning it influences multiple biological systems through regulation of inflammatory signaling, metabolic adaptation, tissue remodeling, and induction of cellular senescence [46]. The cyclin D1 (*CCND1*) gene, widely recognized for its role in cell cycle regulation and interactions with growth factor signaling (e.g., PI3K-AKT, mTOR) and inflammatory pathways (e.g., MAPK), was also targeted by four of the co-expressed miRs identified in this sample [47]. The only gene that ranked above *TGFB1* and *CCND1* in miR targeting was MYC proto-oncogene (*MYC*), a master transcriptional regulator that coordinates cellular growth, proliferation, metabolism, and apoptosis, and amplifies expression of active genes [48]. Collectively, convergent targeting of these mRNAs suggests SASP- and senescence-associated biology may represent the molecular landscape underlying midlife women’s risk for multimorbid depression and T2D. While experimental validation is needed to further clarify the role of senescence- and SASP-related biology, existing pharmacological agents (e.g., metformin, glucagon like peptide-1, senolytics) have shown promise in reducing processes associated with senescent cell accumulation in mixed-sex samples, including modulation of SASP-associated inflammatory signaling and adipose tissue remodeling [49–51]. Future studies may evaluate the preventative benefits of targeting senescence biology in midlife women.

In addition to mechanisms associated with stress/biological weathering and inflammation, findings also revealed regulatory networks that signal involvement of metabolic and endocrine processes. *MYC, CCND1, TGFB1*, vascular endothelial growth factor A (*VEGFA*), and insulin-like growth factor 1 receptor (*IGF1R*) are endocrine-sensitive and involved in metabolic regulation, as well as inflammatory processes [45,48,53–55]. *VEGFA* and *IGF1R* also regulate vascular permeability [54–56], which has potential implications for depression risk related to alterations in the blood brain barrier and common T2D complications (e.g., retinopathy, nephropathy, cardiovascular disease). Overall, these findings suggest that women’s risk for multimorbid depression and T2D in midlife is likely not isolated to a single system or pathway. Rather, interacting processes across biological domains (i.e., metabolic, endocrine, inflammatory, stress-related processes) may act synergistically, contributing to latent progression of multisystem dysregulation that heightens risk for multimorbidity that largely unrecognized within current screening paradigms.

### Limitations

The mRNA targets and enriched pathways were derived from *in silico* analyses; however, we applied the most stringent filters for rigor of human data from miRTarBase to limit the potential for false positives. This dataset lacked quantitative information about menopausal status. Physiological changes related to reproductive aging occur along a continuum spanning multiple years and vary in timing across individuals [8]. By focusing on the midlife timeframe (ages 40-64 years), we increase the potential to capture mechanisms operating across this continuum, including those that may vary in magnitude or direction across different stages of reproductive aging. Depressive symptoms were assessed using the BDI-I, which limits comparisons with other measures and the revised BDI (BDI-II) that is the current version of the instrument. Although significantly higher in the high-risk phenotype, scores were in the minimal to mild range (0-9 minimal; 10-18 mild) [26], likely due to DPP exclusion criteria. However, subclinical depressive symptoms commonly contribute to functional impairments that adversely affect sustained engagement with health promoting behaviors (i.e., exercise, dietary changes, sleep) [57].

## CONCLUSION

Leveraging co-expressed miRs previously shown to be associated with high psychometabolic risk in midlife women, this study extends our prior work by revealing predicted mRNAs and related biological pathways that may underly risk for multimorbid depression and T2D in this high-risk population. The mechanistic findings presented in this manuscript suggest that depression and T2D co-occurrence in midlife women may reflect syndemic clustering, potentially emerging from interactions among metabolic, inflammatory, endocrine, and stress/biological weathering-related processes. Identifying mechanisms underlying risk for multimorbid depression and T2D can inform clinically scalable approaches to enhance risk stratification and guide more effective strategies for prevention and early intervention in this high-risk population. Future research is needed to validate the identified mechanisms and clarify their potential clinical utility.

## Abbreviations

T2D: Type 2 Diabetes
miRs: MicroRNAs
mRNA: Messenger RNA
DPP: Diabetes Prevention Program
KEGG: Kyoto Encyclopedia of Genes and Genomes
SWAN: Study of Women’s Health Across the Nation
FBG: Fasting Blood Glucose
HbA1c: Hemoglobin A1c
BMI: Body Mass Index
HDL: High-density lipoprotein
BDI-I: Beck Depression Inventory
PCA: Principal Component Analysis
RT-PCR: Reverse transcription polymerase chain reaction
P13K-AKT: Phosphatidylinositol 3-Kinase/Protein Kinase B
AGE-RAGE: signaling pathway in diabetic complications:
AMPK: AMP-activated protein kinase
FoxO: Forkhead box O
mTOR: mechanistic target of rapamycin
JAK-STAT: Janus-activated kinase-signal transducers and activators of transcription
*TGF-β*: Transforming growth factor-beta
MAPK: Mitogen-activated protein kinase
Wnt: Wingless-related integration site
HIF: Hypoxia-inducible factor
Rap1: Ras-proximate-1 or Ras-related protein 1
Hippo: Salvador-Warts-Hippo
p53: Tumor protein 53
SASP: Senescence-associated secretory phenotype
*TGFβ1*: Transforming Growth Factor Beta 1
MAPK: Mitogen-activated protein kinase
*MYC*: MYC proto-oncogene, bHLH transcription factor
*CCND1*: cyclin D1
*VEGFA*: *v*ascular endothelial growth factor A
*IGF1R*: insulin-like growth factor 1 receptor

## DECLARATIONS

### Ethics approval and consent to participate

Not applicable.

### Consent for publication

All authors have consented to the publication of this manuscript.

### Availability of data and materials

Data from the DPP trial are available through the NIDDK data repository. MicroRNA data generated from this sample are available from https://doi.org/10.7910/DVN/MJDBJE.

### Authors’ contributions

**KDL:** conceptualization, methodology, formal analysis, visualization, writing - original draft, and writing – review & editing; **BMS**: methodology, conducting the formal analysis, and writing – review & editing. **MG, KAL, SW, NP**: writing – review & editing. **EF**: Supervision of conceptualization, methodology, and formal analysis; writing - original draft, and writing – review & editing

